# Diversity and evolution of centromere repeats in the maize genome

**DOI:** 10.1101/005058

**Authors:** Paul Bilinski, Kevin Distor, Jose Gutierrez-Lopez, Gabriela Mendoza Mendoza, Jinghua Shi, R. Kelly Dawe, Jeffrey Ross-Ibarra

## Abstract

Centromere repeats are found in most eukaryotes and play a critical role in kinetochore formation. Though CentC repeats exhibit considerable diversity both within and among species, little is understood about the mechanisms that drive centromere repeat evolution. Here, we use maize as a model to investigate how a complex history involving polyploidy, fractionation, and recent domestication has impacted the diversity of the maize CentC repeat. We first validate the existence of long tandem arrays of repeats in maize and other taxa in the genus *Zea*. Although we find considerable sequence diversity among CentC copies genome-wide, genetic similarity among repeats is highest within these arrays, suggesting that tandem duplications are the primary mechanism for the generation of new copies. Genetic clustering analyses identify similar sequences among distant repeats, and simulations suggest that this pattern may be due to homoplasious mutation. Although the two ancestral subgenomes of maize have contributed nearly equal numbers of centromeres, our analysis shows that the vast majority of all CentC repeats derive from one of the parental genomes. Finally, by comparing maize with its wild progenitor teosinte, we find that the abundance of CentC has decreased through domestication while the pericentromeric repeat Cent4 has drastically increased.

## Introduction

In spite of the rapid growth in the number of sequenced genomes, centromeres remain poorly understood and relatively cryptic due to their highly repetitive content. Centromere repeats are highly diverse across taxa and their turnover appears to be very rapid (Melters et al, 2013). However, little is known about the genetic mechanisms that produce centromere repeat diversity, though we can begin investigating potential mechanisms with improved assemblies of centromeric regions. Domesticated maize (*Zea mays* ssp. *mays*) has a high quality genome assembly (Schnable et al, 2009) including complete sequence of two centromeres (Wolfgruber et al, 2009), and the breadth of research into maize centromeres makes it one of the best systems to investigate the processes governing centromere repeat evolution.

Maize centromeres are comprised primarily of the 156bp satellite repeat CentC and the CRM family of retrotransposons. Both repeats interact with kinetochore proteins such as CENH3 (Wolfgruber et al, 2009; Zhong et al, 2002) and show variation in abundance across taxa (Albert et al, 2010). While considerable effort has gone to investigating the molecular function of maize centromere repeats (Ananiev et al, 1998; Nagaki et al, 2003; Wolfgruber et al, 2009), we know comparatively little about the evolution responsible for producing the current sequences. CRM elements are better understood, including the age and insertion preferences of different CRM families (Wolfgruber et al, 2009; Sharma et al, 2008). In contrast, no in-depth characterization of the genetic diversity of centromere repeats in the maize genome exists.

In this paper, we describe the patterns of diversity of centromere repeats across the maize genome. We investigate whether the differential ancestry of maize centromeres (Wang and Bennetzen, 2012) has led to chromosome-specific variation of CentC similar to that seen in other species (Kawabe and Nasuda, 2005; Pontes et al, 2004) and how genetic relatedness among individual CentC repeats varies spatially across the genome. We find that CentC copies do not form genetic groups consistent with ancient whole genome duplications or chromosome specificity, despite most large arrays of CentC originating from one of the ancestral subgenomes of maize. We show higher genetic similarity of CentC repeats within clusters, indicating the predominance of tandem duplications in the formation of new CentC copies. Lastly, we use low coverage sequencing and cytological data to show that domesticated maize has less CentC than its wild relatives.

## Methods

### CentC Repeat Identification and Diversity

We downloaded 218 previously annotated CentC sequences (Ananiev et al, 1998; Nagaki et al, 2003) from Genbank. We then searched the maize genome (5b60, www.maizesequence.org) with megaBLAST (McGinnis and Madden, 2004) using the 218 annotated CentCs as a reference, keeping hits with a length of over 140bp and a minimum bit score of 100. After meeting the bit score threshold, the longest hit was retained. We defined CentC’s as being in tandem if the CentC’s start location was within 1000bp of the start location of another CentC.

All 12,162 CentC sequences were aligned using 7 iterations of Muscle (Edgar, 2004) with default parameters. A Jukes-Cantor distance matrix of all sequences was calculated with PHYLIP ((Felsenstein, 1989) http://evolution.genetics.washington.edu/phylip.html), and an unrooted neighbor joining tree was built based on the distance matrix.

We used principle coordinate analysis (PCoA) to cluster CentC variants based on their genetic distances. Eigenvalues from the PCoA were used to determine the number of statistically significant clusters using the Tracy-Widom distribution (Patterson et al, 2006).

We employed the software SpaGeDi ((Hardy and Vekemans, 2002) http://ebe.ulb.ac.be/ebe/Software.html) to estimate the spatial autocorrelation of sequence similarity of CentC repeats in the completely sequenced centromeres 2 and 5. We calculated Moran’s I statistic using Jukes-Cantor genetic distance and measures of physical distance between CentC repeats in base pairs. Confidence intervals for the values of I were estimated by 20,000 random permutations of the physical distances.

Statistical analyses were performed in R with the packages ape (Paradis et al, 2004) and RMTstat (Perry et al, 2009). We compared clusters to chromosome of origin and syntenic maps of maize ancient tetraploidy (Schnable et al, 2011) to determine if the genetic history of maize left a footprint on CentC similarity.

### Read Mapping and Genome Size Correction

We mapped Illumina reads from a broad panel of *Zea* species (Chia et al, 2012; Tenaillon et al, 2011) to a reference consisting of the full complement of 12,162 CentC variants identified in the B73 genome. We also used previously published whole genome chromatin immunoprecipitation (ChIP) (Wolfgruber et al, 2009; Wang et al, 2013) with CenH3. Reads were mapped using Mosaik v1.0 (https://code.google.com/p/mosaik-aligner/). We first optimized mapping parameters by relaxing mapping stringency and evaluating the number of successfully mapped reads with each combination. Consistent with parameters from previous studies mapping reads to repetitive elements (Tenaillon et al, 2011), we required homology to remain at a minimum of 80%. For other non-default parameters, we permuted over many values of hash size, alignment candidate threshold, percent of read aligning, and maximum number of hash positions per seed to find a combination that produced believable alignments. We selected an optimum combination of parameters just below the parameters where we observed a large increase in the total number of reads aligning (Figure S2). Our final set of parameters for tandem repeats used an initial hash size of 8, an alignment candidate threshold of 15 bases, 20% percent of mismatching bases, a minimum of 30% overlap to the reference, and stored the top 100 hits for alignment. After reads were mapped, we calculated the percentage of total reads hitting the given reference and multiplied this value by the relative genome size of each accession as reported in Chia et al (2012) and Tenaillon et al (2011). The total number of reads mapping did not change drastically when using one random copy of CentC versus the full AGPv2 reference, suggesting that our parameters are sufficiently broad to capture genome-wide CentC abundances. Because library preparation has an effect on estimates of repeat abundance (see results), we only used individuals from maize HapMap v2 (Chia et al, 2012) with libraries prepared using identical methods.

We used a different set of mapping parameters for long repeats such as transposable elements. Previous studies (Schnable et al, 2009) estimated that approximately 85% of maize genome derives from transposable elements. Using the short read libraries from Tenaillon et al (2011), we selected parameters so that approximately 85% of the library mapped to the maize transposable element database (www.maizetedb.org) with a minimum homology of 80%. The final parameters for TEs were a hash size of 10, alignment candidate threshold of 11, 80% homology excluding non-aligned portions of the read, and a 30% minimum overlap.

We designed a simulation to estimate the accuracy of our measurements of CentC content (code available at: https://github.com/kddistor/dnasims). In short, our simulations altered the copy number of CentC repeats over a region of fixed length (10Mb), changing the percentage of the genome deriving from the repeat. Illumina reads were simulated from each of the DNA strings and mapped using our pipeline. These simulations showed that our pipeline captured relative differences in abundance well, but underestimated total abundance of CentC. We found that our pipeline could accurately capture differences of 0.05% change in CentC abundance, suggesting that larger differences are likely to be biologically real (Figure S5).

### Simulation of homoplasious mutations

In order to better understand patterns of diversity at CentC, we performed simulations to test the likelihood of homoplasious mutations (i.e. independent mutations occurring at the same position in two different CentC repeats). Our simulation (code available at: https://github.com/paulbilinski/CentC_Analyses/tree/master/Diversity_sims) assumed that CentC has been evolving for 1 million years since the divergence of maize and *Tripsacum* (Ross-Ibarra et al, 2009), a closely related genus whose centromere repeat shares a large amount of homology (Melters et al, 2013). We assumed a constant copy number, a mutation rate of 3×10^−8^ (Clark et al, 2005), and one generation per year.

### PacBio Sequencing

Library preparation and sequencing was performed according to the methods cited in (Melters et al, 2013). Using those protocols, we sequenced one individual from *mays*, *mexicana*, *parviglumis*, and *Z. luxurians* with Pacific Biosciences (Pacific Biosciences, Menlo Park, CA) technology. Approximately 200Mbp of reads were produced from each cell, and reads with length greater than 600bp were retained for analysis of tandem CentC content using BLAST (Table S2). CentC copies were considered tandem if the read had 4 CentC copies within 300bp of each other.

### Fish

Fluorescent in situ hybridization (FISH) was carried out as described in Kato et al (2004) (*Z. luxurians* hybrid) and Shi et al (2010) (*Z. parviglumis* hybrid).

## Results

### Centromere repeats in the maize genome

We found a total of 12,162 CentC copies in the maize reference genome and unassembled BACs. Of these, 8,259 were unique over their full length. No CentC sequence occurred more than 10 times in the genome, and the vast majority (*>*75%, Table S1) of non-unique CentC variants occurred only twice. Of the 2,266 non-unique CentC sequences, only 3 were tandem, identical duplicates. Genome-wide CentC locations also show that nearly all of the 10,639 CentC copies on chromosomes 1-10 are found in clusters; only 14 occurred as solo copies. Clusters varied in width from single CentC copies to 84KB with a mean of *∼*7KB (*∼*45 CentC copies). Chromosomes varied greatly in CentC copy number, though we know that centromere assemblies for all of the chromosomes are not complete. For example, CENH3-ChIP sequence from an oat-maize addition line with one maize chromosome (Kynast et al, 2001) has many reads that map to the unassembled BACs (Table S3). In particular, chromosome 6 has many more reads aligning to the unassembled BACs than it did to its own centromere repeats, suggesting a particularly incomplete assembly. Examining total repeat number, chromosome 7 had the most CentC, with 3,200 copies, while chromosome 6 had the fewest with 32 copies.

We used long-read Pacific Biosciences sequencing to verify that most CentC is in tandem arrays. We sequenced whole genome (*∼* 0.1X) libraries from 4 *Zea* species. In spite of the low coverage, we recovered reads containing CentC sequence from all four taxa (Table S2). In one 6.7KB read from the maize reference line B73, for example, we identified approximately 40 independent CentC copies in tandem, and similar arrays were seen in all four *Zea* species analyzed. These results show that overall structure of the repeats has been maintained for the approximately 140,000 years since the *luxurians*-*mays* divergence (Hanson et al, 1996; Ross-Ibarra et al, 2009) and that a majority of CentC is found in tandem arrays (Table S2).

We then identified how many large clusters of CentC were retained from each of the two parental genomes that comprise the extant maize genome, referred to here as subgenome 1 and subgenome 2 (Figure 1). Previous work identified the parental genome for individual chromosomal segments (Schnable et al, 2011) and centromeres (Wang and Bennetzen, 2012). Because large clusters are less likely to be misassembled, we focused our analyses on the 52 clusters *>*10KB in length (Supplementary Fig S1). We assign clusters to a subgenome if they are flanked by two regions identified as originating from the same subgenome. Thirty-eight of these clusters could be assigned to subgenome 1 (out of 43 assignable). If we restrict the analysis to clusters *>*20KB with clear assignment to one subgenome, all 16 clusters were found in subgenome 1 regions. Even correcting for the genome-wide overrepresentation of subgenome 1 (62.5% of assigned base pairs), these results suggest a strong inequality in the origin of large CentC clusters (Fisher’s exact test, *p <* 0.005 for both 10KB and 20KB clusters). One cluster *>*20KB falls within an unassigned region on chromosome 3.

**Fig. 1.**
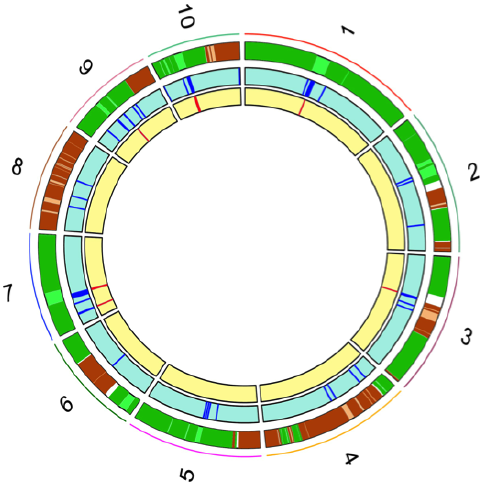
CentC repeat location in relation to the maize subgenomes. The outer ring depicts chromosomal assignment to the two subgenomes, with higher confidence regions are colored with darker colors. Green corresponds to subgenome 1, brown to subgenome 2. Breakpoints between the subgenomes remain uncolored to indicate uncertainty. The middle ring, shaded in blue, displays the locations of all CentCs across the genome. The inner ring, shaded in yellow, displays the locations of all CentC clusters greater than 20KB in length

Previous studies have also described the Cent4 repeat, a tandem pericentromeric repeat that occurs primarily on chromosome 4 (Page et al, 2001). Available evidence does not point to any centromere function for Cent4: CenH3 chromatin immunoprecipitation data (Wolfgruber et al, 2009; Jin et al, 2004) show no significant over-representation of Cent4 compared to five known non-centromeric TE’s, fiber FISH shows clear separation of Cent4 from centromeric repeats (Jin et al, 2004) and Cent4 probes lag behind CentC probes in cell division, suggesting that they are not found in the kinetichore (Jiang et al, 2002; Jin et al, 2004). BLAST analyses of Cent4 sequences from Genbank revealed high homology to the poorly characterized LTR retrotransposon RLX sela that was previously shown to be associated with heterochromatic knobs (Tenaillon et al, 2011; Chia et al, 2012). Cent4 lacks any of the protein sequences necessary for autonomous transposition, such as GAG and POL complexes. But while previous work in rice has documented the presence of nonautonomous LTR retrotransposons in or near the centromere (Jiang et al, 2002), RLX sela also appears to be missing the necessary primer binding sites that would distinguish it as a nonautonomous TE, suggesting that it may be a TE-derived tandem repeat unique to the pericentromere of chromosome 4.

### Relatedness of CentC in the maize genome

CentC copies in the maize genome exhibit tremendous diversity: the overall pairwise identity in our alignment was only 65%, and *∼*98% of sites in the alignment had at least 2 variants. Such diversity led us to ask whether genetic groups of CentC variants could be distinguished. We performed principle coordinate analyses (PCoA) from a genetic distance matrix estimated from our alignment, and assigned individual repeats to genetic clusters following the approach of Patterson et al (2006). We found 58 significant clusters, but observed no pattern of groupings that revealed chromosome specificity of CentC’s or the impact of historical tetraploidy (Figure 2; Table S4).

**Fig. 2.**
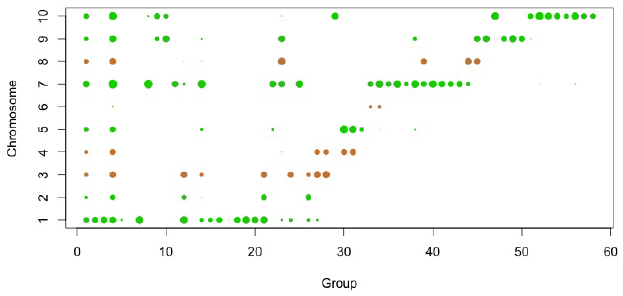
Presence of CentC in each of the heirarchical groups. The 58 clusters found to be statistically significant in forming genetic groups are represented on the x-axis and chromosome of origin on the y-axis. The size of each point is proportional to the log number of sequences in that group on that chromosome. CentC counts from chromosomes whose centromeres were derived from subgenome 1 are colored green and those from subgenome 2 are colored brown

The tandem nature of CentC suggests it increases in copy number through local duplications that produce initially identical copies. Tandem duplication predicts that clusters of CentC should be more closely related than CentC from different clusters. Comparisons of genetic and physical distance among CentC repeats on chromosomes 2 and 5 shows that average genetic similarity is highest within clusters (Figure 3), revealing significant spatial autocorrelation of CentC variants over distances up to 10-50KB (Figure S3 and S4).

**Fig. 3.**
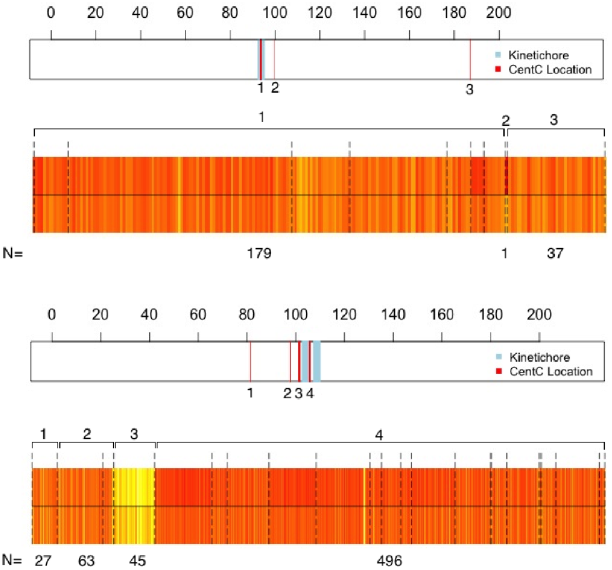
CentC physical location and genetic relatedness for (a) chromosome 2 and (b) chromosome 5. On the physical map, red lines show locations of numbered CentC clusters and blue blocks show the location of the active kinetochores. Scale bar is in MB. Below each physical map is shown a heatmap of genetic relatedness of each CentC to (top row) other copies within its island of tandem repeats delineated by dotted lines and (bottom row) all other copies on the chromosome. Darker colors indicate higher relatedness. The total number of CentC in each cluster is shown below the map

The decreased genetic distance among CentCs in local clusters on chromosome 2 and 5 suggest that many of the genetic groupings discovered in our genome-wide analysis should correspond to local clusters of repeats. However, repeats within individual clusters are frequently found in different genetic groups as defined by principle coordinate analysis (Figure 2). A comparison of shared mutations across all pairs of CentC sequences reveals a potential explanation. Of the *∼* 74 million possible pairs, approximately 6 million share *≥* 2 mutations different from the genome-wide consensus, causing CentC copies to group with sequences that share mutations irrespective of their physical distance. Comparing several triplets at random from our alignment confirms that two sequences in one PCoA assignment share greater pairwise identity than two sequences adjacent to one another in different PCoA groups. A simple forward simulation (see Methods) suggests this pattern could be due entirely to homoplasy rather than long-distance movement of CentC repeats. By stochastically applying mutations to an initially homogeneous group of repeat sequences, we find that plausible parameter values produce *∼* 10 million pairs of repeats sharing *≥* 2 mutations.

### Variation of CentC abundance in *Zea*

Shotgun sequence data from the maize HapMap v2 (Chia et al, 2012), reveals a significantly greater abundance of CentC in teosinte than in domesticated maize (p*<* 0.01; Figure 4). Further support for differences between maize and its wild relatives comes from shotgun sequence data from *Z. luxurians* (Tenaillon et al, 2011). Analysis of these data find nearly twice as much CentC in *Z. luxurians* as the maize inbred B73. To corroborate these results, we performed fluorescent in-situ hybridization of F1 crosses between inbred maize and teosinte to determine if cytological observations agreed with our sequencing findings. FISH data supports our observation that the teosintes *parviglumis* and *Z. luxurians* have more CentC than inbred maize (Figure 4). Using whole genome shotgun PacBio long reads, we further investigated the overall structure of repeats across the different *Zea* species. Percentages of the libraries showing tandem repeats did not differ greatly across the taxa (Figure S2).

**Fig. 4.**
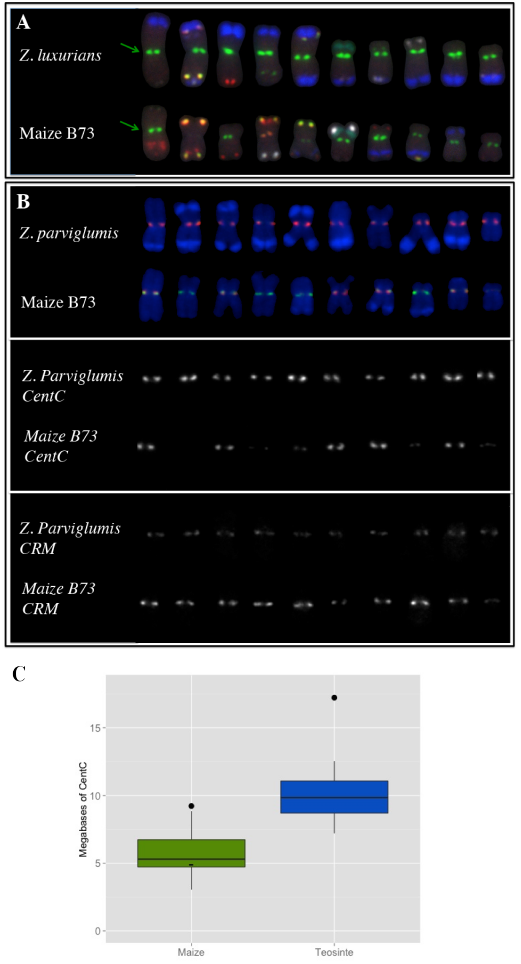
(a) FISH analysis of a single individual heterozygous for B73 and *Zea luxurians* (GRIN accession PI422162). Chromosomes, ordered from 1 (left) to 10 (right), were hybridized with the Birchler probe cocktail (Kato et al, 2004). Green shows CentC and a 4-12-1 subtelomere repeat, blue the 180bp knob repeat, red the abundant TAG microsatellite and another subtelomeric repeat, white the TR1 knob repeat, orange the Cent4 repeat, yellow the 5S rDNA repeat, and aqua shows the Nucleolus Organizer Region. The green signals at the primary constrictions (arrows) are CentC. Note that *Z. luxurians* has far brighter CentC signals. This image was graciously provided by Patrice Albert. (b) FISH analysis of a single individual heterozygous for B73 and *Z. parviglumis*(GRIN accession PI566687). Chromosomes were hybridized with the Shi et al (2010) probe cocktail showing CentC in red and CRM2 in green. Each separate probe is shown separately below the two-color image, highlighting that CentC is more abundant in *Z. parviglumis* and CRM2 is more abundant in maize. (c) Mb of CentC in genomic libraries of maize and teosinte. Box plots show data from Chia et al (2012). Points show data for maize inbred B73 and the teosinte *Z. luxurians* from Tenaillon et al (2011). For comparison, the data point of maize inbred B73 in Chia et al (2012) is shown with a tick mark on the box plot.

## Discussion

Our analysis of centromere repeat diversity across the maize genome identifies thousands of copies exhibiting tremendous diversity. But while we can cluster the repeats into groups of related sequences, these groups have little relation to current or ancient maize chromosomes (Figure 2). We find no evidence of chromosome specific repeats as observed in *Arabidopsis* species (Kawabe and Nasuda, 2005; Pontes et al, 2004), suggesting the presence of a mechanism that homogenizes repeats across centromeres on different chromosomes. We further show that Cent4, once thought to be a chromosome-specific centromere repeat (Page et al, 2001), appears to be a poorly characterized tandem repeat or nonautonomous retroelement, but is not associated with the centromere. Inter-chromosomal transposition was postulated by Shi et al (2010) as an explanation for marker genotypes showing evidence of recombination, and such transposition could also explain the similarity of repeats across all chromosomes.

Additionally, we find that virtually all the large arrays of CentC in the maize genome derive from one of the two ancestral genomes present in modern day *Zea* (Figure 1). One possible explanation for the biased inheritance pattern is that centromere inactivation mirrors the differential expression seen between the subgenomes. Centromere inactivation could then lead to differential loss of CentC clusters, such that large clusters remain only in the active centromeres. Large CentC clusters are biased in favor of subgenome 1, the subgenome known to have lower gene loss and higher average expression than maize subgenome 2 (Schnable et al, 2011). Consistent with a correlation between expression and CentC loss in subgenome 2, Reinhart and Bartel (2002) found evidence of regulatory microRNA’s corresponding to centromeric repeats. An alternative explanation is that the subgenome 1 parent had more or larger CentC arrays, leading to an initial unequal representation in the ancient polyploid ancestor of maize.

Our sequence comparison of CentCs also enabled us to explore the relationship between genetic and physical distance among repeats. Using the well-assembled centromeres on chromosomes 2 and 5, we found spatial autocorrelation of relatedness among repeats, and we observe genome-wide that many CentC’s within an array fall into the same genetic cluster. Our observations are consistent with the simple idea that most repeats arise due to tandem duplication, and long distance transposition of CentC, while necessary to homogenize repeats across chromosomes (Shi et al, 2010), is relatively uncommon.

One unusual result from our sequence comparison was the finding that pairs of CentC on different chromosomes share very high sequence similarity. Our simulations suggest that, under realistic assumptions about mutation rate and divergence time, such a pattern is possible due to homoplasious mutation alone. Roughly 80% of the CentC repeats have their closest genetic relative on the same chromosome, as expected under a model of tandem duplication, but only 14% of closest genetic pairs are found within 10KB of each other. We speculate that the vast majority of CentC’s in the genome are thus a result of relatively old tandem duplications, and that sufficient time has occurred since duplication for rearrangements and mutations to break up patterns of identical tandem repeats.

Previous cytogenetic work identified differences in centromere repeat content between domesticated maize and its wild relatives *Z. mays* ssp. *parviglumis*, *Z. mays* ssp. *mexicana*, and *Z. luxurians* (hereafter *parviglumis*, *mexicana,* and *luxurians*) (Albert et al, 2010) but was unable to quantify differences. Our resequencing results show that while there is little difference in the distribution of CentC in tandem arrays, the absolute abundance of CentC has decreased during domestication, and we verify this with FISH in two maize-teosinte hybrid individuals (Figure 4). Variability in observed abundance of transposable elements (Chia et al, 2012) suggests that the decrease seen in CentC is not due to causes common to all repetitive sequences. The maize genome is smaller than its teosinte counterpart, largely due to differences in the abundance of heterochromatic knobs (Poggio et al, 1998). Zhang and Dawe (2012) have postulated an adaptive relationship between centromere size and genome size based on an observed correlation between centromere size and genome size across a number of grass species. A model correlating centromere size to total genome size would propose that the decrease in CentC abundance seen post-domestication is due to selection for smaller active centromeres to complement the smaller overall genome size. While our current data are insufficient to evaluate this conclusion, future work investigating differences in CentC content among maize landraces that vary in genome size (Poggio et al, 1998) may provide an opportunity to further test our hypothesis.

In conclusion, our detailed study of centromere repeats in the maize genome has highlighted differential contribution of subgenome, spatial autocorrelation along a chromosomes, and changes in abundance over the short time scale of domestication. Further work evaluating CentC evolution across multiple populations and multiple related species may shed additional light on the timing and causes of these changes.

## Acknowledgements

We wish to thank Pacific Biosciences for sequencing resources. We thank Patrice Albert and James Birchler for providing the high-quality FISH image of the *Z. luxurians* X B73 hybrid. We thank Siddharth Bhadra-Lobo, Vince Buffalo, Anne Lorant, Gernot Presting, Lauren Sagara, Michelle Stitzer, and National Science Foundation summer exchange program interns Cesar Alvarez Mejia, Aurelio Hernandez Bautista for advice and helpful discussion. This project was funded by National Science Foundation grant IOS-0922703.

## Supplemental Material

**Fig. S1.**
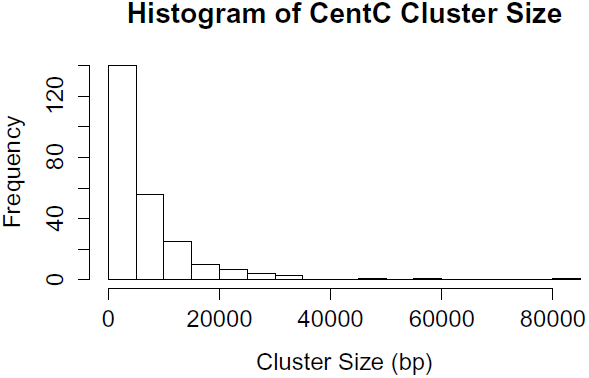
CentC cluster size across all chromosomes.

**Fig. S2.**
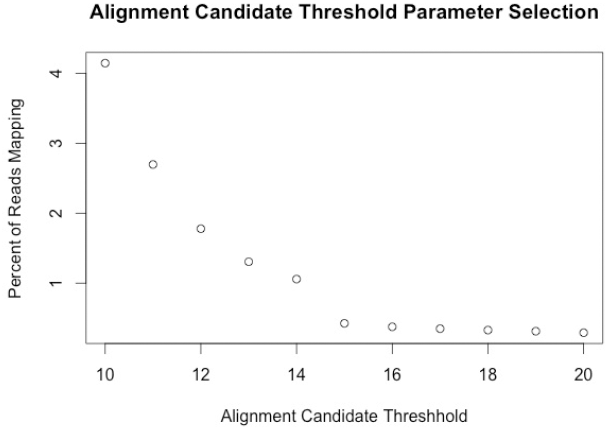
Parameter selection for Alignment Candidate Threshold (ACT) for Mosaik. All other parameters were kept constant while ACT was changed. ACT was the parameter for which a non-linear pattern was observed. We selected to use an ACT of 15, the value for which we observed the greatest relative decrease between total percent mapping values. The sharp change suggests that, at a lower ACT, we may be mapping a non-CentC element to our reference.

**Fig. S3.**
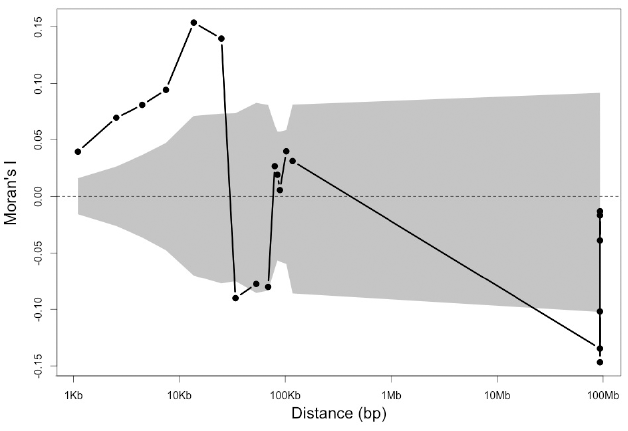
Measure of Moran’s I for Chromosome 2. Gray areas show the confidence interval, calculated using permutations of genetic distance.

**Fig. S4.**
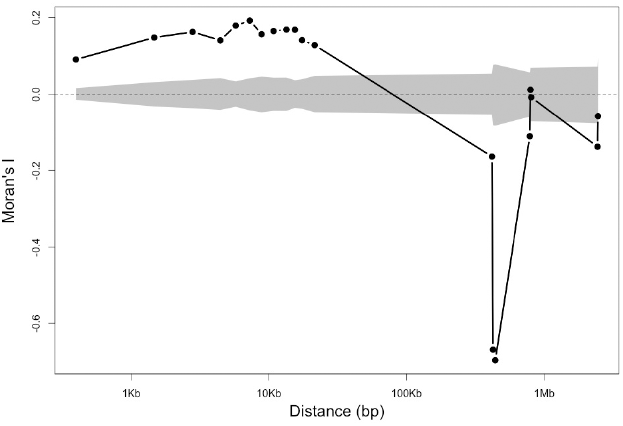
Measure of Moran’s I for Chromosome 5. Gray areas show the confidence interval, calculated using permutations of genetic distance.

**Fig. S5.**
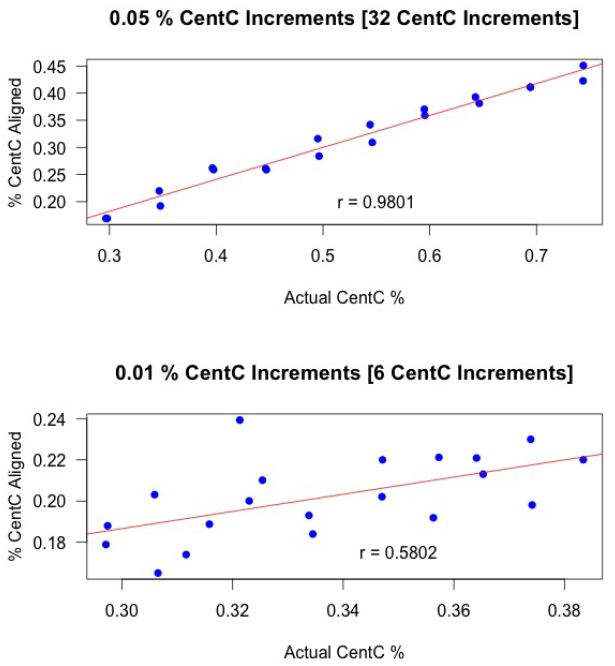
Graphs showing our ability to capture changes in CentC repeat abundance under constant genome size. We simulated 10MB of DNA with varying CentC content and simulated Illumina reads from the DNA. Reads were mapped with our Mosaik pipeiline, and several simulations at each percentage of genomic content were performed.

**Table S1.** CentC Occurence Count In Maize RefGenv2

| Occurrences | Number of CentC's |
| --- | --- |
| 1 | 8259 |
| 2 | 1233 |
| 3 | 263 |
| 4 | 89 |
| 5 | 31 |
| 6 | 13 |
| 7 | 7 |
| 8 | 0 |
| 9 | 0 |
| 10 | 1 |

**Table S2.** PacBio Read Counts and Tandem CentC

| Maize Line | Reads over 600bp | Reads with $\geq 4$ CentC | % Reads Showing Tandem CentC |
| --- | --- | --- | --- |
| B73 | 237995 | 30 | 0.030252736 |
| *luxurians* | 156964 | 79 | 0.050330012 |
| *mexicana* | 141939 | 150 | 0.1056792 |
| *parviglumis* | 227050 | 89 | 0.039198414 |

**Table S3.** ChIP Reads mapping to Unassembled from different Oat-Maize Addition (OMA) Lines. Reads were mapped using Bowtie2 (Langmead and Salzberg, 2012) with the paramater-very sensitive-local.

| File Key | Maize Chr | Percent Reads Aligning to Unassembled BACs |
| --- | --- | --- |
| JJ1BU (OMA 6.34) | 6 | 21.35 |
| JJ1BR (OMA 1.36) | 1 | 15.56 |
| JJ1CF (OMA 9.41) | 9 | 6.16 |
| JJ1CH (OMA 8.05) | 8 | 6.69 |
| JJ1CG (OMA 10.26) | 10 | 9.21 |
| JJ1CI (OMA 8.05) | 8 | 8.5 |

**Table S4.**
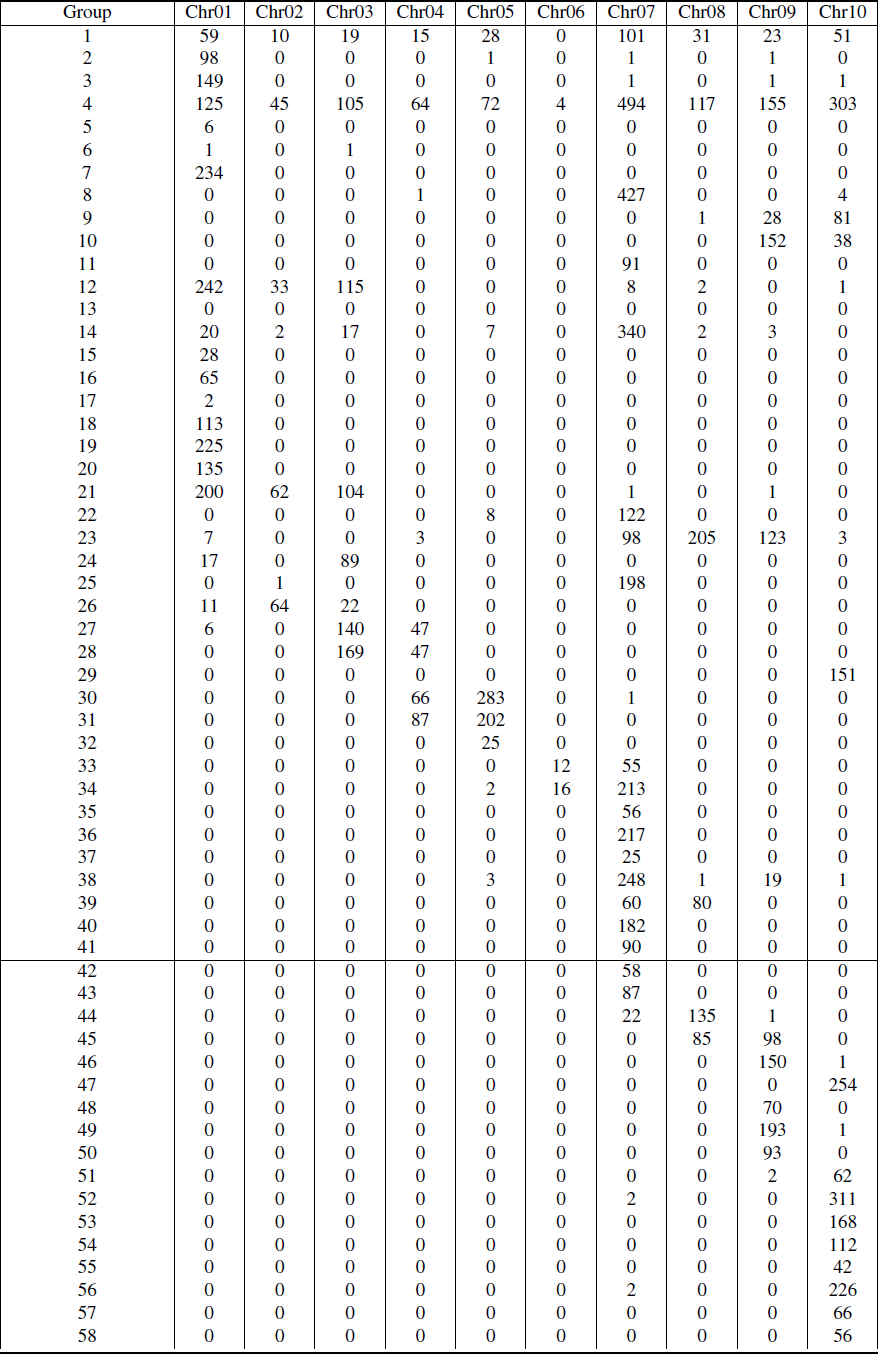
Heirarchical clustering group assignment for copies of CentC, sorted by chromosome. The number of CentC’s from each chromosome is represented in the table.

